# Impacts of the Samarco tailing dam collapse on metals and arsenic concentration in freshwater fish muscle from Doce River, southeastern Brazil

**DOI:** 10.1101/2020.05.09.086058

**Authors:** Frederico Fernandes Ferreira, Mariella Bontempo Duca de Freitas, Neucir Szinwelski, Natállia Vicente, Laila Carine Campos Medeiros, Carlos Ernesto Gonçalves Reynaud Schaefer, Jorge Abdala Dergam, Carlos Frankl Sperber

## Abstract

On November 2015, Samarco tailings dam in Mariana MG, Brazil, collapsed, releasing 62 million tons of tailings that advanced through 668 km of the Doce River and adjacent floodplain. Although being the worst environmental disaster in Brazil, little is known about consequences to aquatic biota. Here we evaluate the effects of the tailings mudflow on metal and arsenic concentration in fish and how concentration correlates with water and fish characteristics. We quantified semitotal amounts of Ag, Al, As, Cd, Cr, Cu, Fe, Hg, Mn, Ni, Pb and Zn in fish muscle tissue using ICP-MS in 255 individuals (34 species) sampled in unaffected and affected along the Doce River basin. Arsenic and Hg were higher in fish from affected sites, likely due to turbulent mixing of previously sedimented material by the giant tailings wave. Silver and Zn concentrations were higher in unaffected sites. Arsenic concentration in *Geophagus brasiliensis* decreased with increasing fish weight. Copper and Zn decreased with increasing fish weight considering the whole assembly of fish. The tailings mudflow increased water conductivity and conductivity increased Al concentration in fish, so we expected a larger Al concentration in fishes from affected sites. However, the observed Al concentration in fishes from affected sites was lower than expected by water conductivity. Thus, the tailings mudflow reduced Al uptake or accumulation in fishes. Mercury decreased with increasing water conductivity in both unaffected and affected sites considering all species and in *G. brasiliensis* alone. Despite the relatively low concentration range of metals and As found in fish, fishes from sites affected by the iron ore tailings mudflow showed higher As and Hg concentration, compared to fishes from unaffected sites. The higher As and Hg in affected sites require further detailed monitoring to ensure safeguards of human health by fishing activity along the Doce River.

## Resumo

Em novembro de 2015, a barragem de rejeitos da Samarco em Mariana, MG, Brasil, rompeu, liberando 62 milhões de toneladas de rejeitos de minério de ferro que avançaram por 668 km do rio Doce e planície adjacente. Embora este tenha sido considerado o pior desastre ambiental do país, pouco se sabe sobre as consequências da liberação do fluxo de lama para a biota aquática. Aqui avaliamos os efeitos do fluxo de lama de rejeitos sobre a bioacumulação de metais e arsênio em várias espécies de peixes coletadas na bacia do rio Doce e como a bioacumulação se correlaciona com as características da água e dos peixes. Quantificamos as quantidades semitotais de Ag, Al, As, Cd, Cr, Cu, Fe, Hg, Mn, Ni, Pb e Zn no tecido muscular de peixes usando ICP-MS em um total de 255 indivíduos (representando 34 espécies), amostrados em locais não afetados e afetados ao longo da bacia do rio Doce. As concentrações de As e Hg foram maiores nos peixes dos locais afetados,, provavelmente devido à mistura turbulenta de material previamente sedimentado pela onda gigante de rejeitos. As concentrações de Ag e Zn foram maiores nos peixes de locais não afetados. A concentração de As na espécie mais abundante nos locais não afetados e afetados (*Geophagus brasiliensis*) diminuiu com o aumento do peso dos peixes. A concentração de Cu e Zn diminuiu com o aumento do peso dos peixes, considerando toda a assembleia de espécies. O fluxo de lama de rejeitos aumentou a condutividade da água e a condutividade aumentou a concentração de Al nos peixes, portanto esperávamos uma maior concentração de Al nos peixes de locais afetados. No entanto, a concentração de Al observada nos peixes de locais afetados foi menor do que o esperado pela condutividade da água. Assim, o fluxo de lama de rejeitos reduziu a assimilação ou o acúmulo de Al nos peixes. O mercúrio diminuiu com o aumento da condutividade da água nos locais não afetados e afetados, considerando todas as espécies e também dentro de *G. brasiliensis*. Apesar da faixa de concentração relativamente baixa de metais nos peixes, os peixes de locais afetados pelo fluxo de lama de rejeitos de minério de ferro apresentaram maior concentração de As e Hg, em comparação com peixes de locais não afetados. A maior concentração de As e Hg nos locais afetados requer um monitoramento mais detalhado para garantir segurança alimentar em relação à atividade pesqueira ao longo do rio Doce.

## INTRODUCTION

The Samarco dam rupture on November 5th, 2015, in Mariana, MG, Brazil, was one of the largest ever reported failures of a tailing dam worldwide (Hatje et al. 2017). The dam collapse released more than 62 million tons of iron ore tailings into the riverbeds and floodplains, reaching the Gualaxo do Norte, Carmo and Doce River, causing the death of 19 people and polluting 668 km of downstream watercourses down to the Atlantic Ocean (Fernandes et al. 2016; dos Santos Vergilio et al. 2020). The Doce River has a long history of land degradation and unplanned water use, with high rates of deforestation followed by erosion, sedimentation, pollution and eutrophication (Pires et al. 2017). Gold mining, intense in the area since the 17th century up to present days, polluted the riverbeds with toxic elements, such as Hg and As (Hatje et al. 2017). Although anthropogenic disturbances, such as agriculture and mining activities have caused long term habitat degradation, the dam collapse differs from these sources of pollution due to the large amount and speed that the tailings’ wave travelled, altering the structure of the habitats, such as alluvial soils, river sediments, terraces, floodplains and river banks (Hatje et al. 2017).

Samarco assured that the tailings waste was composed mainly of inert mineral particles (Escobar 2015). However, several metals and As were detected in the Doce River after the disaster, despite no previous monitoring was available for assessing the process and time of contamination (Carvalho et al. 2017, Hatje et al. 2017). This can be potentially harmful for human health and aquatic biota. Here we aimed evaluating if the discharge of the iron ore tailings, produced by the collapse of Samarco dam in Mariana, MG, Brazil, altered the concentration of selected metals and As in fishes from the Doce River basin.

## METHODS

### Sampling sites and determination of metals and arsenic concentration in fish

We sampled 12 sites within the Doce River basin (Figure 1) from November 2018 to March 2019 (three years after the Samarco dam rupture). Fish samples and water characteristics were collected in Piranga (sites 01 and 02), Piracicaba (site 03) and Santo Antônio (sites 04, 05, 06 and 07) rivers, areas not reached by the tailings mudflow and considered unaffected sites for this study (Figure 1). The affected sampling sites included Gualaxo do Norte (site 08), Carmo (site 09) and Doce (site 10, 11 and 12) rivers (Figure 1). In each sampling site we set 20 fishing nets of 10 m2, comprising ten different mesh sizes, ranging from 15 to 80 mm knot to knot distance. Fish nets were set at sunset and were collected in the next morning, after 12 hours. Each captured fish was anesthetized with clove oil (Eugenol), identified, weighted (g), measured (mm), sexed, and maturation status of each fish’s gonad was evaluated macroscopically, and each specimen was photographed. We extracted a sample of ca. 20g of muscle tissue of each specimen using ceramic knifes sterilized with 10% nitric acid. All packaging material and dissection tools, including plastic cutting board, were sterilized before and after using them at each sampling site. All captured fish specimens were packaged in separate plastic bags for each species in the sample, duly labeled with taxonomic classification, sampling site and collection date, for reference collection, deposited at the Ichthyological Collection of the João Moojen Zoology Museum of the Federal University of Viçosa (UFV). Fish were fixed in 10% formaldehyde solution for a period of 48 hours and then transferred to 70% ethanol solution. Fish sampling, euthanasia and transport was authorized by the Biodiversity Information and Authorization System – SISBIO, Chico Mendes Institute for Biodiversity Conservation – ICMBio, Ministry of Environment – MMA (SISBIO authorization number 55430-2). One muscle tissue sample of each fish specimen was sent to the Tommasi Ambiental Environment and Food Analysis laboratory, based in Vitória, ES, for quantitative determination of metals and arsenic. In the laboratory, fish samples (~1.5 g) were crushed and digested in a Microwave Multiwave GO (Anton-Paar, Austria) using a mixture of 65% nitric acid and 30% hydrogen peroxide. The resulting solution was transferred quantitatively to a polypropylene tube and the volume completed with type I water at 10 mL and subsequently analyzed by ICP-MS. Water parameters (pH, O2 and conductivity) were recorded in each site, using an Akso-AK88 (Akso ©) multi-parameter.

**Figure 1.**
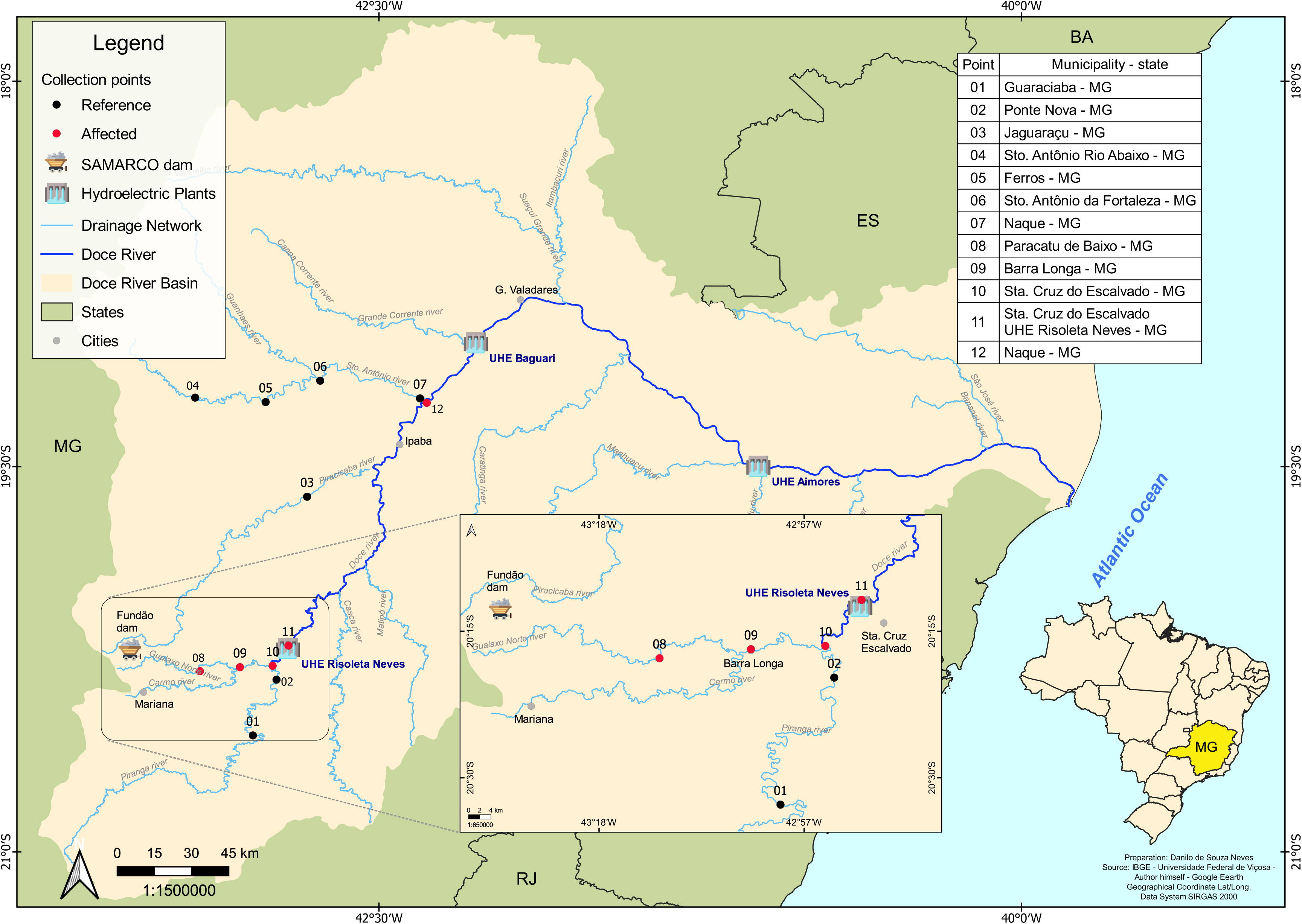
Map of the Doce River basin (in light rose), in Minas Gerais and Espirito Santo states, Brazil. Sampling sites are represented as black (unaffected) and red (affected) circles.

### Statistical analyses

Response variables were log-transformed to reduce residual asymmetry. All statistical models used normal distribution and were run under R (R Core Team 2019). We used two statistical approaches: multivariate and univariate. The multivariate approach included two- and three-way multivariate analyses of variance (MANOVAs), both with all fish species combined and with a subset of the collected fish species, including only data on the fish species collected in sufficient abundance in both unaffected and affected sites (*Geophagus brasiliensis*). To deal with the hierarchical structure of our data, in which fish individuals were nested within sites, we included ‘site’ as explanatory factor, analogous to block, in the MANOVAs (Snijders and Bosker 2011). We used the elements as response variables in the MANOVAs, to evaluate the effects on fishes elements profile. When significant effects of the of the tailings mudflow in the MANOVAs were detected, we investigated which element concentrations were significantly different comparing unaffected and affected sites using univariate mixed effects models (one-way analysis of variance (ANOVAs) and analyses of covariance (ANCOVAs), considering each element concentration as the univariate response variable. We present significant results of ANOVAs as mean + standard deviation; significant results for ANCOVAs are present as scatterplots of log-transformed observed values and lines for values predicted by the minimum adequate models. We used the tailings mudflow as a two-level explanatory factor, and water parameters (pH, O2, conductivity) and fish characteristics (weight and sex) were co-variables, together with the interaction terms between fish characteristics and mudflow. The univariate models were adjusted as generalized linear models with mixed effects (GLMMs), with site as random intercept (Gelman and Hill 2006). Through this approach the hierarchical structure of our data was fully dealt with (Zuur et al. 2009). Univariate ANOVAs and ANCOVAs where run for all fish species combined, disregarding the fish species, and then again only for *Geophagus brasiliensis*. Further details on the statistical analyses are available at Supplemental Data (SD) Methodology Details, including data transformations and the equations of the multi- and univariate models, and further references. Indication of which fish species were included in each analysis are available at SD Table S1.

## RESULTS

We sampled 255 adult fish belonging to 34 species (Table 1). Two hundred and two individuals, belonging to 26 species, were collected in unaffected sites; 53 individuals, belonging to 21 species, were collected in affected sites; 13 species were present in both unaffected and affected sites. The profile of the element concentrations in fish muscle tissue was significantly different between unaffected and affected sites (Two-way MANOVA; P<2.2^−16^; SD Table S2). There was significant interaction of fish species with the effect of the tailings mudflow, considering all species combined (Three-way MANOVA; P=0.014; SD Table S3), as well as a significant effect of fish species (P<2^−16^) and tailings mudflow (P<2^−16^) affecting the profile of the elements concentration in fish. The results found for the analysis when species were combined agreed entirely with the results that we found considering only the six fish species that presented at least three individuals in unaffected and affected sites (SD Table S4). For two of these six species (*G. brasiliensis* and *P. adspersus*) we detected an effect of the tailing mudflow on the profile of the elements concentrations (SD Table S5), but only *G. brasiliensis* was collected in sufficient number of individuals at both sites to enable further analyses.

**Table 1.** Number of fish individuals collected per species in sites unaffected (n=7) and affected (n=5) by the Samarco iron ore tailings dam rupture, in the Doce River basin, ordered according to decreasing accumulated abundance, and number of sites where the species occurred. Information on the species included in each of the statistical analyses are available in Supplemental Data: Table S1.

| sp # | Fish species | Unaffected sites | Affected sites | Total | Number of sites |
| --- | --- | --- | --- | --- | --- |
| 1 | *Geophagus brasiliensis* | 30 | 6 | 36 | 9 |
| 2 | *Oligosarcus argenteus* | 27 | 1 | 28 | 8 |
| 3 | *Astyanax lacustris* | 21 | 0 | 21 | 6 |
| 4 | *Hoplias intermedius* | 10 | 7 | 17 | 8 |
| 5 | *Hypostomus affinis* | 13 | 4 | 17 | 6 |
| 6 | *Pachyurus adspersus* | 16 | 1 | 17 | 5 |
| 7 | *Loricariichthys castaneus* | 14 | 2 | 16 | 4 |
| 8 | *Megaleporinus conirostris* | 9 | 4 | 13 | 6 |
| 9 | *Astyanax sp. 2* | 10 | 0 | 10 | 2 |
| 10 | *Hypomasticus mormyrops* | 10 | 0 | 10 | 2 |
| 11 | *Pimelodus maculatus* | 3 | 4 | 7 | 3 |
| 12 | *Oreochromis niloticus* | 4 | 3 | 7 | 4 |
| 13 | *Henochilus wheatlandii* | 7 | 0 | 7 | 2 |
| 14 | *Hoplias malabaricus* | 1 | 5 | 6 | 2 |
| 15 | *Rhamdia quelen* | 2 | 3 | 5 | 3 |
| 16 | *Hypostomus auroguttatus* | 5 | 0 | 5 | 4 |
| 17 | *Cichla piquiti* | 4 | 0 | 4 | 1 |
| 18 | *Leporinus copelandii* | 4 | 0 | 4 | 3 |
| 19 | *Salminus brasiliensis* | 0 | 3 | 3 | 1 |
| 20 | *Prochilodus vimboides* | 0 | 2 | 2 | 1 |
| 21 | *Lophiosilurus alexandri* | 1 | 1 | 2 | 2 |
| 22 | *Trachelyopterus striatulus* | 1 | 1 | 2 | 1 |
| 23 | *Astyanax fasciatus* | 2 | 0 | 2 | 2 |
| 24 | *Crenicichla lacustris* | 2 | 0 | 2 | 2 |
| 25 | *Cyphocharax gilbert* | 2 | 0 | 2 | 1 |
| 26 | *Delturus carinotus* | 2 | 0 | 2 | 2 |
| 27 | *Astyanax sp. 1* | 0 | 1 | 1 | 1 |
| 28 | *Hoplosternum littorale* | 0 | 1 | 1 | 1 |
| 29 | *Leporinus macrocephalus* | 0 | 1 | 1 | 1 |
| 30 | *Myloplus asterias* | 0 | 1 | 1 | 1 |
| 31 | *Prochilodus costatus* | 0 | 1 | 1 | 1 |
| 32 | *Prochilodus lineatus* | 0 | 1 | 1 | 1 |
| 33 | *Gymnotus carapo* | 1 | 0 | 1 | 1 |
| 34 | *Pseudauchenipterus affinis* | 1 | 0 | 1 | 1 |
| **Total** |  | **202** | **53** | **255** | **12** |

### Arsenic and Mercury concentrations

The concentration of As and Hg in fish muscle tissue were higher in affected compared to unaffected sites (As: P=0.00018; Hg: P=0.016; Figures 2A-B; SD Table S6). When we included water and fish characteristics as co-variables (see SD Table 9 for water parameter values), we detected significant interaction of co-variables with the effects of the tailing mudflow for As (fish sex*tailings mudflow: P=0.018; fish weight*tailings mudflow: P=3.69-6; SD Table S7). In the unaffected sites, As concentration increased with fish weight (P< 2.2^−16^; Figure 3A). In affected sites, there was no effect of fish weight on As concentration (P=0.35), but males showed higher As concentration than females (P=0.0027; Figure 3A). For *G. brasiliensis*, however, As decreased with increasing fish weight (P=0.0078; Figure 3D; SD Table S8), was higher in affected sites (P=8.24-4) and showed no differences between gender (P=0.64).

**Figure 2.**
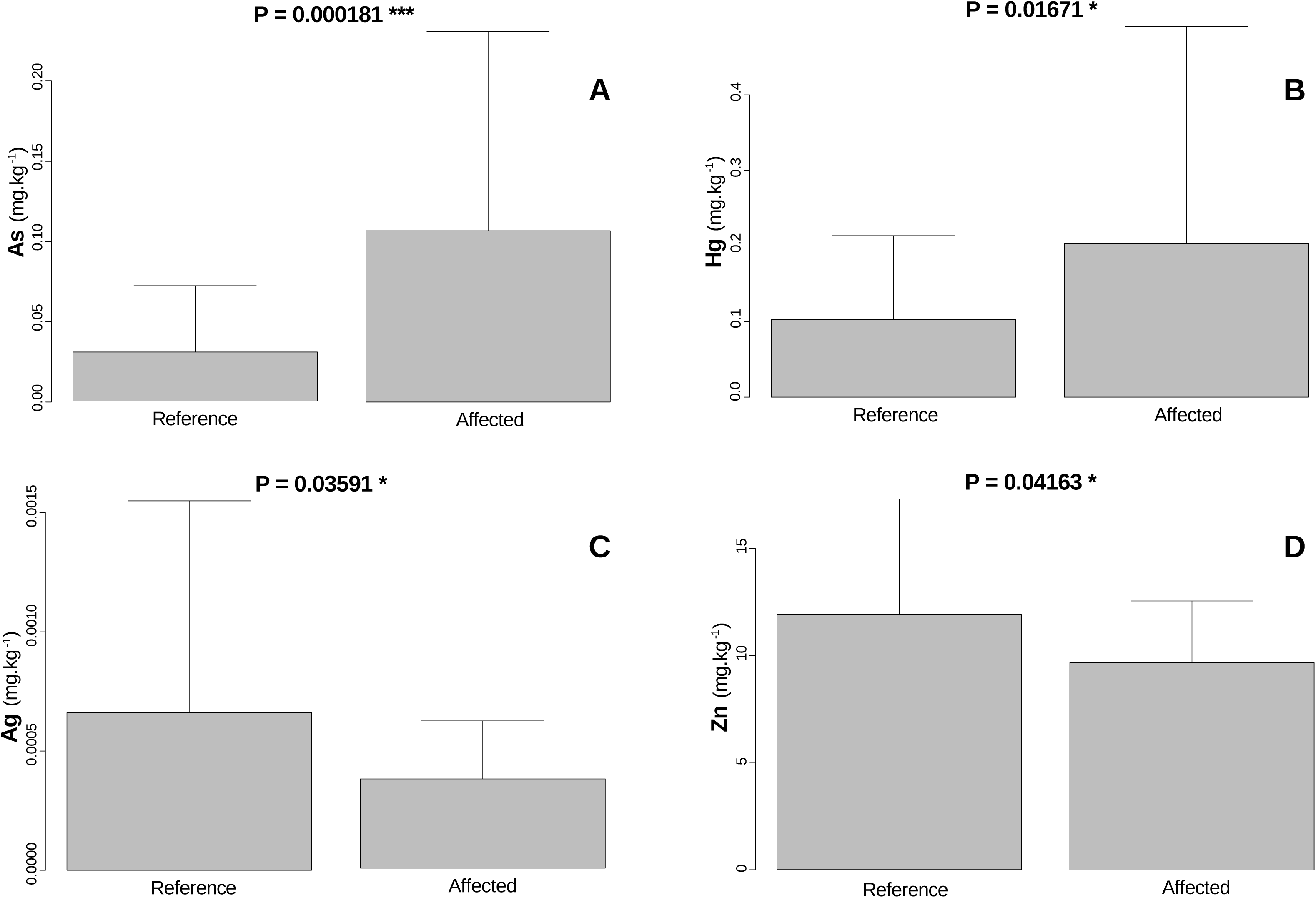
Metals and arsenic concentrations (Means ± SD) in fish muscle tissue (mg.kg^-1^) of all species combined; (A) arsenic(As), (B) mercury (Hg), (C) silver (Ag), and (D) zinc (Zn) in sites that were unaffected or affected by the iron ore tailings mudflow that resulted from the Samarco dam rupture. P-values refer to mixed effects ANOVAs (GLMMs), with site as random intercept. Only significant results (P<0.05) are presented.

**Figure 3.**
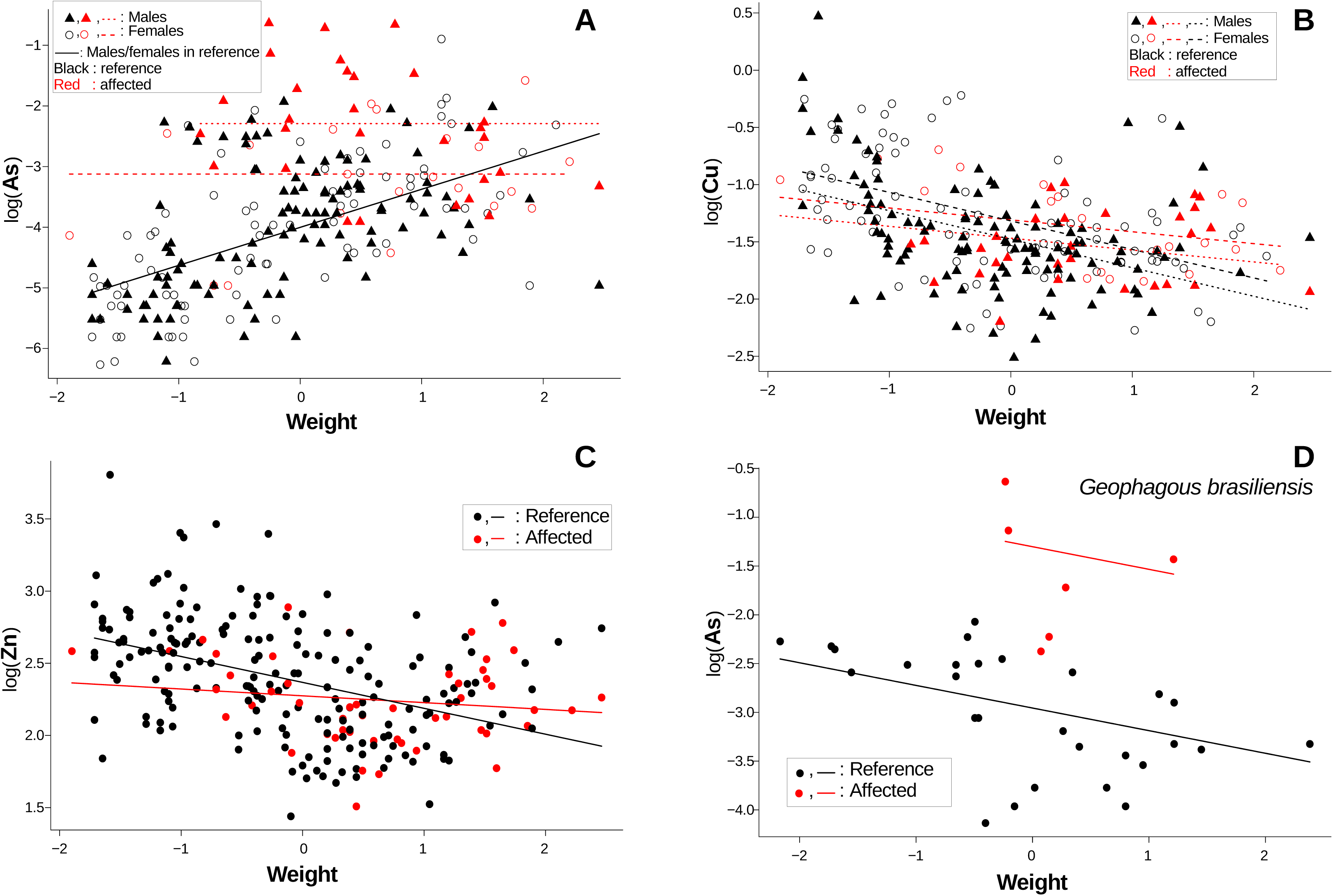
Concentration (log transformed) in fish muscle tissue, in relation to fish weight (centered and scaled), considering the whole fish species assembly, of (A) arsenic (As), (B) copper (Cu) and (C) zinc (Zn); and (D) arsenic (As) within *Geophagous brasiliensis*, comparing fishes from unaffected (black) and from affected (red) sites. Lines refer to predicted values for the minimum adequate mixed effects model (ANCOVA Generalized Linear Mixed Effects Model), with site as random intercept. Lines parallel to the horizontal axis mean that fish weight did not affect the response variable; when there are parallel lines (the red lines in (A), same color lines in (B), red and black lines in (D), this means that there was no interaction of the categorical explanatory variables – sex in (A) and (B), unaffected vs. affected sites in (D), with fish weight, meaning that the effect of weight was the same between fish sexes (A, B) or between unaffected and affected sites (D). Different lines parallel to each other mean significant effect of the categorical variable (sex in(A) and (B), unaffected vs. affected sites in (D)). (A) As increased with fish weight in unaffected sites, but not in affected sites, and As was higher in males than in females in affected sites; (B) Cu decreased with fish weight in both unaffected and affected sites, but this decrease was steeper in unaffected than in affected sites, and Cu was higher in females than males in both affected and unaffected sites; (C) Zn decreased with fish weight in unaffected, but not in affected sites; (D) As in *Geophagous brasiliensis* decreased with fish weight in both unaffected and affected sites and was higher in afected than unaffected sites.

Mercury concentration in fish decreased with increasing water conductivity (P=0.0052; Figure 4B) and was higher in affected than in unaffected sites (Figure 4B). Male fishes showed lower Hg concentration than females (Figure 4B). For *G. brasiliensi*s Hg concentration also decreased with increasing water conductivity (P=0.0015; Figure 4C) and was higher in affected sites (P=8.33^−5^; SD Table S8).

**Figure 4.**
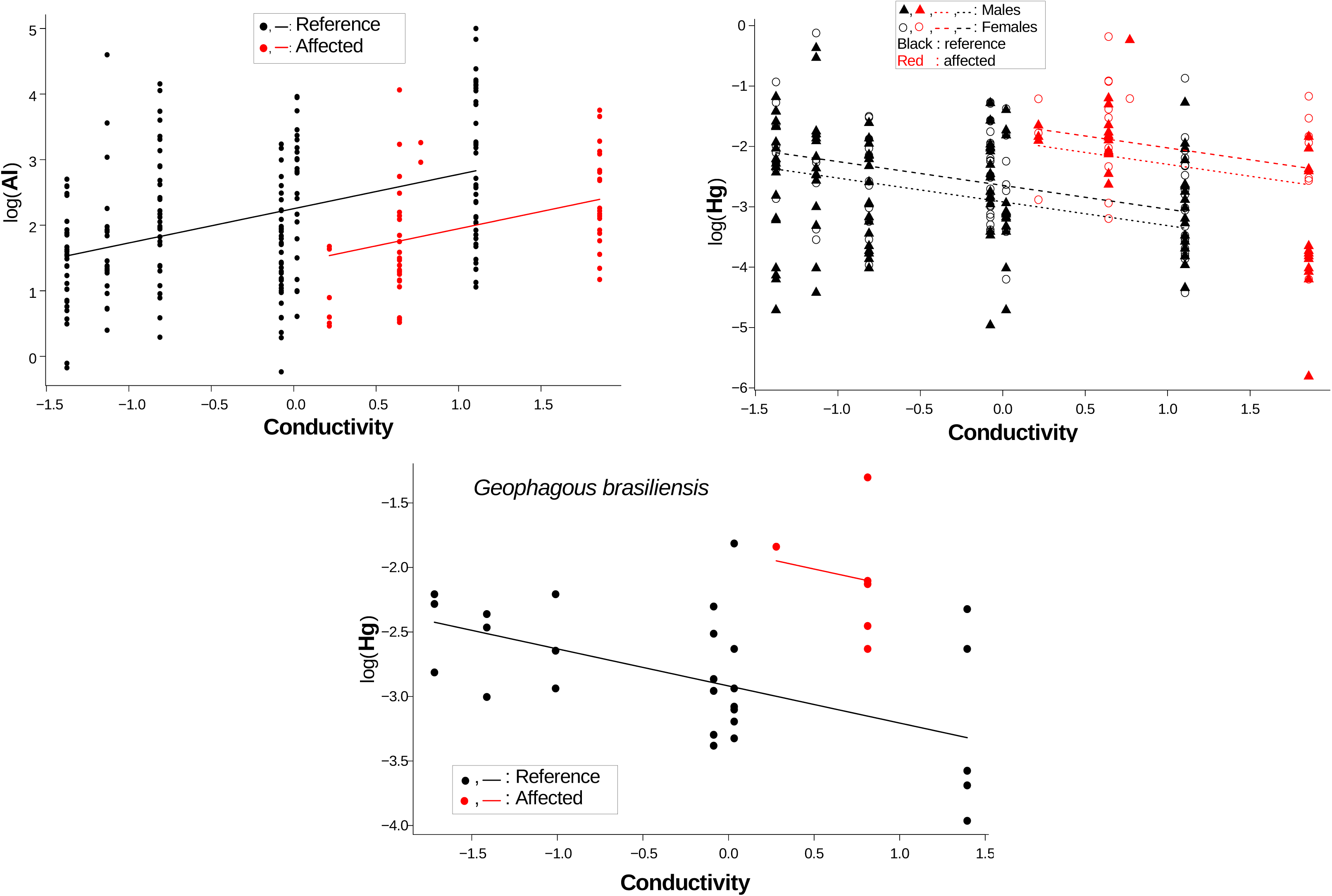
Concentration (log transformed), in fish muscle tissue, in relation to water conductivity, considering the whole fish species assembly, of (A) aluminum (Al) and (B) mercury (Hg); and of (C) mercury (Hg) within *Geophagous brasiliensis*, comparing fishes from unaffected (black) and from affected (red) sites. Lines refer to predicted values for the minimum adequate mixed effects model (ANCOVA Generalized Linear Mixed Effects Model), with site as random intercept. (A) Al increased with water conductivity and was lower in affected than in unaffected sites; (B) Hg decreased with watr conductivity, was higher in affected sites, and was higher in females than in males; (C) Hg in *Geophagous brasiliensis* decreased with water conductivity and was higher in affected than in unaffected sites.

### Silver and Zinc concentrations

Silver (P=0.035) and zinc (P=0.041) concentrations in fish muscle tissue were lower in affected than in unaffected sites (Figure 2C-D; SD Table S6). Zinc concentration decreased with increasing fish weight (P=0.031; SD Table S7) (Figure 3C), but this decrease was less evident in affected sites. When we included water and fish characteristics as co-variables to explain Ag concentration, the effects of the iron tailings mudflow disappeared (P=0.81; SD Table S7).

### Aluminum and Copper concentrations

Aluminum and Cu concentrations did not differ between unaffected and affected sites (Al: P=0.86; Cu: P=0.79; SD Table S6), but when we included water and fish characteristics as co-variables, there was a significant effect of the tailings mudflow on Al (P=0.017; SD Table S7) and a significant interaction of tailing mudflow with fish weight on Cu concentration (P=0.044; SD Table S7). Cu concentrations decreased with increasing fish weight (Figure 3B), but this decrease was less steep in affected sites (Figure 3B). Copper concentration was lower in male fishes compared to females (Figure 3B). Aluminum concentration increased with increasing water conductivity (P=0.0033; Figure 4A), and was lower in affected sites (P=0.017).

### Cadmium, Chrome, Iron, Manganese, Nickel and Lead concentrations

Cadmium, Cr, Fe, Mn, Ni and Pb did not differ between unaffected and affected sites (P>0.14; SD Table S6). Including water parameters and fish characteristics as co-variables did not reveal any effect of the tailings mudflow on these metals concentration (P>0.14; SD Table S7).

## DISCUSSION

To our knowledge, this is the first evaluation of the effects of the Samarco iron ore dam collapse on the concentration of metals in tissues of freshwater fish from the affected area. Our results demonstrate that As and Hg concentration in fish was higher in affected than unaffected sites. Segura et al. (2016) verified that the tailings mud did not contain high levels of As nor Hg, leading us to discard the hypothesis that the tailing mud itself was the source of the increased levels of these elements in fish tissue from the affected sites. Arsenic has a regional geological affinity with gold bearing sulphides (Au in arsenopirite) (Alloway 1990), whereas Hg was used in gold extraction procedures since 17th century, mainly in the upper Doce river tributaries, like Carmo and Gualaxo (Schaefer et al. 2015). This shows the historical transference of both elements to Doce River. The disturbance of the riverbed sediments caused by the passage of the tailings wave provoked mixing and uplifting of old riverbed sediments exposing otherwise unavailable and stabilized elements (Lambertz and Dergam 2015). The tailing spill caused a substantial increase in suspended sediment loads and Hg increase in the sediments, whereas As increased in suspended particulate matter (Hatje et al. 2017). Our results are a strong evidence that the disturbance of the river bed, through the tailings mudflow, made As and Hg bioavailable in the affected sites, leading to larger As and Hg concentration in the fishes from these sites. Increased As and Hg concentrations in all fish species from affected, compared to unaffected sites, are in agreement with the intraspecific results for *G. brasiliensis*, supporting the consistency of these results. The observed increase in As and Hg concentrations in fish from affected sites might be further augmented in the long term, when enhanced erosion caused by heavy rain episodes may lead to remobilization and transport of contaminated particles to the affected sites (Hatje et al. 2017).

Zinc concentration was lower in fishes of affected, compared to unaffected sites, which shows that the tailing mudflow reduced the bioavailability of Zn or the tailings mudflow washed the former bioavailable Zn in the affected sites. The tailing mud flow might also have diluted Zn concentration in the water or the interference of the tailings mudflow on water parameters might have reduced Zn uptake by fishes, altering chemical processes of metal elements’ availability (Başyiğit and Tekin-Özan 2013) and absorption by the fishes (Niyogi and Wood 2004).

Once the increase of Al uptake with water conductivity was taken into account, we unveiled that Al uptake was lower in affected sites (Fig. 4A). Thus, although the alterations in water parameters caused by the tailings mudflow (increased water conductivity) should increase Al concentration (as observed in the unaffected sites), there was a further, yet unknown, alteration in affected sites, that reduced Al uptake or accumulation in fishes. This reduction in Al concentration countered also the increased dissolved Al levels in affected compared to unaffected sites detected by Hatje et al. (2017). Thus, our results indicate that there was a physical-chemical or biological alteration, provoked by the tailings mudflow, that countered the Al bioavailability processes related to water conductivity, and countered the larger dissolved Al brought by the tailings mud. Further studies, on fish physiology and Al bioaccumulation, are necessary to unveil these processes.

### Relationship of As, Cu and Zn with fish weight

Our results show evidence of As bioaccumulation with weight in unaffected sites, but not in affected sites, where this element showed higher levels of concentration in fish muscle tissue irrespective of fish weight, showing that in affected sites the bioaccumulation process with fish weight was absent (Fig. 3A). This result might be an artifact of the changes of As concentration in caused by the tailings mudflow and the short-term contact of fishes with this new environmental conditions. If this is actually artifact, this pattern should change, once the continuous exposure to As eventually results in its bioaccumulation with fish weight (Kumari et al. 2017). On the other hand, our results might reveal a change in fish uptake or accumulation of As in affected sites. Another important aspect of this result is the chronic As contamination revealed in our study through the bioaccumulation of this element in unaffected sites, which highlights the historical mismanagement of water resources in these areas.

Considering our results for *G. brasiliensis*, As concentration decreased with increasing body weight in both unaffected and affected sites. This finding suggests that, in this species, the excretion of As overcomes its uptake. Fish have evolved different mechanisms for biotransformation of As to less toxic forms, which are then readily excreted (Bears et al. 2006). Geophagy is frequently related to As exposure in primates (Krishnamani et al. 2000), and it is likely that it plays a role also in this fish species, which is a bottom feeder (Abelha et al. 2004). An efficient As excretion would have a high adaptive value for a species that ingests high levels of toxic compounds. Thus, decreased levels of As in fish with higher body weights might be a specific adaptation for these *G. brasiliensis’* populations. Further studies would be necessary to evaluate this hypothesis.

In contrast to As, Cu and Zn concentration decreased with increasing fish weight in both sites, which may be an evidence of some sort of biodilution. Yohannes et al. (2013) verified Cu interspecific biodilution in fish. Biodilution is commonly associated with an interspecific process along the food web, but it may also occur intraspecifically, as when it results from fast growth rates (Pickhardt et al. 2002). For both Cu and Zn, the decrease with fish weight was less steep in affected than unaffected sites, indicating a convergent reduction in the bioaccumulation of these elements in the sites affected by the tailings mudflow. Goethite (FeO(OH)), abundant in iron ore tailings (Weissenborn et al. 1994), adsorbs Cu (Christophi and Axe 2014). Thus, the tailings mud might have reduced these elements’ bioavailability in the affected sites through adsorption by goethite.

## CONCLUSIONS

Despite the relatively low concentration range, river waters affected by the iron ore tailings mudflow, provoked by the Samarco iron ore dam rupture, showed higher As and Hg concentration in fish muscle tissue. The tailings mudflow also decreased water conductivity and altered As, Al, Cu and Zn bioaccumulation in fish, showing that environmental assessment of metal and As elements in biota should include water chemistry, weight and sex of fish as co-variables. The higher Hg and As in affected sites is require further detailed monitoring to ensure safeguards of human health by fishing activity along the Doce river. Long-term studies should include testing for synergistic effects, as well as for effects of water chemistry on bioavailability and bioaccumulation, in order to provide a sound and safe understanding on element effects on the biota and human food safety.

## Supporting information

SD Supplemental Data Details on the Statistical analyses

SD Table S1: Fish species, number of individuals, sites and indication for which statistical analyses they were included

SD Table S2: Results of the two-way MANOVA for all fish species

SD Table S3: Results of the three-way MANOVA for all fish species

SD Table S4: Results of the two-way MANOVA for the six fish species that presented at least three individuals in each affected and reference sites

SD Table S5: Results of the two or three-way MANOVAs

SD Table S6: Mean concentration of metals and arsenic in fish from sites unaffected and affected and results of the mixed effects one-way ANOVAs

SD Table S7: Results of the mixed effects ANCOVAs

SD Table S8: Results of the mixed effects ANCOVAs for Geophagous brasiliensis

SD Table S9: Mean + standard deviation (SD) of water parameters

## REFERENCES

Abelha MCF, Goulart E. 2004. Oportunismo trófico de Geophagus brasiliensis (Quoy & Gaimard, 1824)(Osteichthyes, Cichlidae) no reservatório de Capivari, Estado do Paraná, Brasil. Acta Scient Biol Sci 26(1): 37–45.

Alloway B J. Heavy Metals in Soils, John Wiley and Sons: New York, 1990.

Bears H, Richards JG, Schulte PM. 2006. Arsenic exposure alters hepatic arsenic species composition and stress-mediated gene expression in the common killifish (Fundulus heteroclitus). Aquat Toxicol 77:257–266.

Başyiğit B, Tekin-Özan S. 2013. Concentrations of some heavy metals in water, sediment, and tissues of pikeperch (Sander lucioperca) from Karataş Lake related to physico-chemical parameters, fish size, and seasons. Pol J Environ Stud 22(3), 633–644.

Carvalho MS, Moreira RM, Ribeiro KD, Almeida AM. 2017. Concentração de metais no rio Doce em Mariana, Minas Gerais, Brasil. Acta Brasiliensis. 1(3): 37–41.

Christophi CA, Axe L. 2014. Competition of Cd, Cu and Pb adsorption on goethite. Journal of Environmental Engineering. 126 (1): 66–74.

dos Santos Vergilio C, Lacerda D, de Oliveira BCV, Sartori E, Campos GM, de Souza Pereira AL, de Rezende CE. 2020. Metal concentrations and biological effects from one of the largest mining disasters in the world (Brumadinho, Minas Gerais, Brazil). Sci Rep 10(1): 1–12.

Escobar H. 2015. Mud tsunami wreaks ecological havoc in Brazil. Science. 350 (6265): 1138–1139.

Fernandes GW, Goulart FF, Ranieri BD, Coelho MS, Dales K, Boesche N, Bustamante M, Carvalho FA, Carvalho DC, Dirzo R et al. 2016. Deep into the mud: ecological and socio-economic impacts of the dam breach in Mariana, Brazil. Natureza & Conservação. 14 (2): 35–45.

Gelman A, Hill J. 2006. Data analysis using regression and multilevel/hierarchical models. Cambridge University Press.

Hatje V, Pedreira RM, de Rezende CE, Schettini CAF, de Souza GC, Marin DC, Hackspacher PC. 2017. The environmental impacts of one of the largest tailings dam failures worldwide. Sci rep 7(1), 1–13.

Kumari B, Kumar V, Sinha AK, Ahsan J, Ghosh AK, Wang H, DeBoeck G. 2017. Toxicology of arsenic in fish and aquatic systems. Environ chem let 15(1), 43–64.

Lambertz M, Dergam J. 2015. Mining disaster: huge species impact. Nature. 528 (7580): 39.

Niyogi S, Wood CM. 2004. Biotic ligand model, a flexible tool for developing site-specific water quality guidelines for metals. Environmental Science and Technology. 38(23): 6177–6192.

Pickhardt PC, Folt CL, Chen CY, Klaue B, Blum JD. 2002. Algal blooms reduce the uptake of toxic methylmercury in freshwater food webs. Proc Nat Acad Sci 99(7): 4419–4423.

Pires AP, Rezende CL, Assad ED, Loyola R, Scarano FR. 2017. Forest restoration can increase the Rio Doce watershed resilience. Persp Ecol Conserv 15(3), 187–193.

R Core Team. 2013. R: A language and environment for statistical computing. Vienna (AT): R Foundation for Statistical Computing. [accessed 2019 Jan 22]. http://www.r-project.org

Segura FR, Nunes EA, Paniz FP, Paulelli ACC, Rodrigues GB, Braga GÚL, Batista BL. 2016. Potential risks of the residue from Samarco’s mine dam burst (Bento Rodrigues, Brazil). Environ Poll 218, 813–825.

Snijders TA, Bosker RJ. 2011. Multilevel analysis: An introduction to basic and advanced multilevel modeling. Sage.

Schaefer CEGR, dos Santos EE, de Souza CM, Neto JD, Fernandes Filho EI, Delpupo C. 2015. Cenário histórico, quadro físiográfico e estratégias para recuperação ambiental de Tecnossolos nas áreas afetadas pelo rompimento da barragem do Fundão, Mariana, MG. Arq Mus Hist Nat Jardim Bot UFMG 24(1-2).

Weissenborn PK, Dunn JG, Warren LJ. 1994. Quantitative thermogravimetric analysis of haematite, goethite and kaolinite in Western Australian iron ores. Thermochim Acta 239, 147–156.

Yohannes YB, Ikenaka Y, Nakayama SM, Saengtienchai A, Watanabe K, Ishizuka M. 2013. Organochlorine pesticides and metals in fish from Lake Awassa, Ethiopia: insights from stable isotope analysis. Chemosphere. 91(6): 857–863.

Zuur A, Ieno EN, Walker N, Saveliev AA, Smith GM. 2009. Mixed effects models and extensions in ecology with R. Springer Science and Business Media.

