## Supplementary material for "Impacts of the Samarco tailing dam collapse on metals and arsenic concentration in freshwater fish muscle from Doce River, southeastern Brazil": SD Supplemental Data Details on the Statistical analyses

For the statistical analyses, we considered only adult fishes. We adjusted concentration of the 12 evaluated elements as response variables. Response variables were log-transformed to reduce residual asymmetry. When concentration values were equal to zero or when they were below the detection level of the ICP-MS, we substituted their values for the minimum detected concentration of that element, before calculating the logarithm, so as to avoid non-existing logs for zero values and, at the same time, to minimize departure from the observed concentrations. All statistical models used normal distribution and were run under R (R Core Team, 2019). We evaluated significance of the fixed explanatory terms by step-wise deletion of non-significant terms, using likelihood ratio (Likelihood Ratio Tests – LRT) using chi-squared distribution (Crawley, 2013). To report non-significant results ($P>0.05$), we aggregated them citing the lowest P value of these results. We used two statistical approaches, one multivariate, another univariate.

***Multivariate analyses to evaluate fishes' element profiles.***

The multivariate approach evaluated if each fish individual's profile of element concentrations was affected. This approach was meant to avoid multiple tests on the same replicate, the fish individual. In the multivariate analyses, each fish individual was considered an independent replicate (n=255). To deal with the hierarchical structure of our data, in which fish individuals were nested within sites, we included 'site' as explanatory factor (Snijders & Bosker, 2012), analogous to blocks. This means that in the multivariate analyses we disregarded the correlation of fishes from the same site. We used Pillai's test to evaluate significance.

***Two-way MANOVA.***

To evaluate the effects of the tailings torrent on the profile of elements' concentrations in the fish muscle tissue, we run a two-way multivariate analysis of variance (two-way MANOVA), with normal distribution. In this two-way MANOVA we adjusted the 12 elements' concentration (log-transformed) as the multivariate response variable (for indication of which species were analyzed, see Two-way MANOVA (eq. 1) in Supplemental Data Table 1). Explanatory terms were 'affected by the tailings torrent' (affected or reference site), as a two-level explanatory factor, and site as a 12-level effect analogous to block, for all fish species altogether, as in the equation below:

$$Y_{i,j} \sim \text{tailings torrent}_k + \text{site}_l + \varepsilon_{i,j} \qquad \text{(eq. 1)}$$

where $Y_{i,j}$: the matrix of the elements' concentrations (log transformed) in the fish muscle tissue for each element (index i: 1 to 12) in each fish individual (index j: 1 to 255)[1]; tailings torrent (index k: 1 or 2)[2]: explanatory two-level factor; site (index l: 1 to 12): block effect; $\varepsilon_{i,j}$: residuals with normal distribution, considering individual fishes as replicates (n=255). A significant effect of the tailings torrent would mean that there were significant differences in the profile of the concentrations of the estimated elements in fish muscle tissue between affected and reference sites. Significant effect of site would mean that there are differences among sites irrespective of the tailings torrent. As far as the aim of this study was to evaluate the effects of the tailings torrent, significant effects of site alone discarded our working hypothesis.

***Three-way MANOVA.***

1 $Y_{i,j}$ represents a matrix, in this case with 12 columns (one column for each element, the index i varying from 1 to 12) and 250 lines (on line for each fish individual, the index j varying from 1 to 250).

2 When a variable has only one index, such as 'tailings torrent$_k$', it represents a vector, in this case with two values, either 1 or 2, corresponding to the two levels of the factor (affected or reference).

To evaluate if the effects of the tailings torrent on the profile of elements' concentrations in the fish muscle tissue, differed among fish species, we run a three-way multivariate analysis of variance (three-way MANOVA), with normal distribution, also with the 12 elements' concentration (log-transformed) as the multivariate response variable, but in which the explanatory terms were 'affected by the tailings torrent' (affected or reference site), as a two-level explanatory factor, 'fish species identity' (a factor with the number of levels equal to the number of collected fish species), the interaction term of 'fish species identity' with 'tailings torrent', and site as a 12-level effect analogous to block. In this three-way MANOVA we included all fish species (for indication of which species were analyzed, see Two-way MANOVA (eq. 1) in Supplemental Data Table 1), as in the equation below:

$$Y_{i,j} \sim \text{tailings torrent}_k + \text{fish species identity}_m + tt*fsi + \text{site}_l + \varepsilon_{i,j} \qquad \text{(eq. 2)}$$

where $Y_{i,j}$: the matrix of the elements' concentrations (log transformed) in the fish muscle tissue for each element (index i: 1 to 12) in each fish individual (index j: 1 to 255); tailings torrent (index k: 1 or 2): explanatory two-level factor; fish species identity (index m: 1 to 34): explanatory factor with 34 levels; tt*fsi: interaction of tailings torrent with fish species identity; site (index l: 1 to 12): block effect; $\varepsilon_{i,j}$: residuals with normal distribution, considering individual fishes as replicates (n=255). If there was a significant interaction of fish species identity with the effects of the tailings torrent, it meant that different fish species responded differently to the tailings torrent. If there was no interaction of fish species identity with the effects of the tailings torrent, this meant that the fish species responded similarly to the tailings torrent. As far as our aim was to evaluate the effects of the tailings torrent, significant effects of fish species or site alone discarded our working hypothesis.

***Strict three-way MANOVA.***

A significant interaction between 'fish species identity' and 'tailings torrent' in the three-way MANOVA above, could mean that the response to the tailings torrent differed among fish species, but it could also be an artifact of heterogeneous number of individuals in each species. Thus, to discard the artifact effect, we run a second three-way MANOVA (we called it 'strict three-way MANOVA), with exactly the same statistical model of eq. 2, but exclusively with those fish species that presented at least three individuals in each affected and reference sites (for indication of which species were analyzed, see Strict three-way MANOVA (eq. 2) in Supplemental Data Table 1). If the results of the first three-way MANOVA were robust, we expected agreement with the results of the strict three-way MANOVA. If the results of the strict three-way MANOVA did not corroborate the results of the first three-way MANOVA, we considered the first results as artifact.

***Intraspecific MANOVAs.***

If there was significant interaction of the effects of fish species identity with the effects of the tailings torrent in the three-way MANOVAs above, we run separate MANOVAs for each fish species, adjusting the estimated elements' concentration (log-transformed) as the multivariate response variable, so as to evaluate if the elements' profile was affected by the tailings torrent in each species. These analyses were only possible for those species that presented sufficiently high number of individuals to avoid residual rank deficiency (for indication of which species were analyzed, see Intraspecific MANOVAs (eq. 2, 3) in Supplemental Data Table 1). Residual rank deficiency occurs when there is insufficient information contained in the data to estimate the model. When there were sufficient number of individuals, we adjusted two-way MANOVAs, with the same explanatory terms as in equation 1, that is, with both 'tailings torrent' and 'site' as explanatory variables. When there were insufficient number of individuals to adjust the two-way MANOVA, we adjusted a one-way MANOVA, disregarding the hierarchical structure of our sampling, as in the equation below:

$$Y_{i,j} \sim \text{tailings torrent}_j + \varepsilon_{i,j} \qquad \text{(eq. 3)}$$

where $Y_{i,j}$: the matrix of the elements' concentrations (log transformed) in the fish muscle tissue of each individual fish within the evaluated species, for each element (index i: 1 to 12), in each fish individual (index j: 1 to 255); tailings torrent (index k: 1 or 2): explanatory two-level factor; $\varepsilon_{i,j}$: residuals with normal distribution, considering individual fishes as replicates. These analyses were run for a subset of fish species (see MANOVAs per species (eq. 2, 3) in Supplemental Data Table 1). The number of replicates depended on the number of collected fish individuals in the analyzed species.

***Univariate mixed-effects models.***

If we found significant effects of tailings torrent in the profile of the concentrations of elements, that is, if the term 'tailings torrent' was significant in the two-way MANOVA (eq. 1) adjusted for all fish species altogether, our next step was to investigate which elements' concentrations were effectively different between affected and reference sites. To answer this question, we adjusted separate univariate statistical models, one for each of the evaluated elements, with each elements' concentration as response variable (log-transformed). These univariate models were adjusted as generalized linear models with mixed effects (GLMMs), with site as random intercept (Gelman & Hill, 2006). Through this approach the hierarchical structure of our data was fully dealt with (Zuur et al., 2009). We evaluated significance of the fixed explanatory terms by step-wise deletion of non-significant terms, using likelihood ratio (Likelihood Ratio Tests – LRT) using chi-squared distribution (Crawley, 2013). These analyses were run for all fish species (for indication of which species were analyzed, see GLMMs ANOVAs and ANCOVAs (eq. 4, 5) in Supplemental Data Table 1).

***Mixed-effects one-way ANOVAs.***

To evaluate which of the elements' concentrations in the fishes' muscle tissue were affected by the tailings torrent, we adjusted mixed-effects one-way ANOVAS, as in the equation below:

$$Y_j \sim \text{tailings torrent}_k + (1|\text{site}_l) + \varepsilon_{j|l} \quad \text{(eq. 4)}$$

where $Y_j$: the vector of each elements' concentration (log-transformed) in the fish muscle tissue of each individual fish (index j: 1 to 240)[3]; tailings torrent (index k: 1 or 2: explanatory two-level factor; $(1|\text{site}_k)$[4]: random intercept; $\varepsilon_{j|l}$: normal distributed residual, nested according to site, considering individual fishes as replicates (n=255). We adjusted separate models for each of the 12 evaluated elements. Significant effect of tailings torrent meant that the concentration of the evaluated element in fish muscle tissue differed between affected and reference sites.

***Mixed-effects ANCOVAs, with water and fish characteristics as co-variables.***

As far as water and fish characteristics might alter uptake and accumulation of elements by fishes, there could be differences among sites and among fishes that affected the elements' concentration in fish muscle tissue. To evaluate and control for that, we adjusted GLMMs with the characteristics of water and of fish as co-variables in the univariate statistical models, with each elements' concentrations as univariate response variable in separate models for each element. In these models, the explanatory variables were the two-level factor 'tailings torrent' (affected or reference site), the five co-variables: water pH, dissolved O2, water conductivity, fish weight and fish sex (also as a two-level factor), together with the two level interactions of fish weight and fish sex with the factor 'tailings torrent'. Significant

3 In the univariate analyses, there is only one response variable per statistical model, thus $Y_j$ is a vector, with so many lines as replicates (individual fishes), indicated by the index j. In these analyses, j varied from 1 to 240. We discarded the data of 15 fishes due to lack of water parameter data.

4 This notation refers to random intercept, given the site, where the vertical bar (|) represents "given".

interactions of fish characteristics with the factor ‘tailings torrent’ would indicate that the effect of the tailings torrent on the element’s concentration was different between male and female fishes, or that the tailings torrent altered the response of the element’s concentration to fish weight. Thus, to evaluate the eventual effects of water and fish characteristics upon the effects of the tailings torrent, we adjusted mixed-effects analyses of co-variance (ANCOVA), as in the equation below:

$$Y_j \sim \text{tailings torrent}_k + \text{water parameters}_{n|k} + \text{fish characteristics}_{o|j} + \text{fish characteristics*tailings torrent} + (1|\text{site}_l) + \varepsilon_{j|l} \quad \text{(eq. 5)}$$

where $Y_j$: the vector of each elements’ concentration (log-transformed) in the fish muscle tissue of each individual fish (index j: 1 to 240); tailings torrent (index k: 1 or 2): explanatory two-level factor; three water parameters (index n: 1 to 3) measured at each site (index k: 1 to 10)[5]: water pH, water conductivity and dissolved O2; fish characteristics (index o: 1 to 2), evaluated for each individual fish (index j): weight and sex[6]; the two level interactions of fish characteristics (weight and sex) with the factor ‘tailings torrent’; $(1|\text{site}_l)$: random intercept; $\varepsilon_{j|l}$: normal distributed residual, nested according to site. We adjusted separate models for each of the 12 evaluated elements. For these analyses we had only 10 sites (n=10), because for two sites we had trouble with the multi-parameter meter in the field. Significant effect of any two-level interaction meant that the effect of the tailings torrent differed between fish sexes or was altered by fish weight. Significant effect of the tailings torrent meant that the concentration of the evaluated element in fish muscle tissue differed between affected and reference sites. As far as our aim was to evaluate the effects of the tailings torrent, significant effects of water parameters or fish characteristics alone would discard our working hypothesis.

5 We measured three water parameters, represented by the index n, being either 1, 2 or 3 (respectively water pH, water conductivity and dissolved O2), in each site (index k). The vertical bar (|) represents “given”, meaning that water parameters are the same for all fish individuals collected in the same site.

6 Fish characteristics (weight and sex) are nested within fish individuals, represented by the vertical bar (|).

***Intraspecific GLMM ANCOVA.***

For *Geophagous brasiliensis*, the most abundant fish species, we were able to adjust a GLMM ANCOVA for the 12 elements, so as to evaluate if within fishes of this species, the elements' concentrations were affected by the tailings torrent or any of the co-variables related to water or fish characteristics (indicated as Intraspecific GLMM ANCOVA (eq. 6) in Supplemental Data Table 1), according to the equation below:

$$Y_j \sim \text{tailings torrent}_k + \text{water parameters}_{n|k} + \text{fish characteristics}_{o|j} + (1|\text{site}_l) + \varepsilon_{j|l} \qquad \text{(eq. 6)}$$

where $Y_j$: the vector of each elements' concentration (log-transformed) in the fish muscle tissue of each individual fish (index j: 1 to 240) within the analyzed species; tailings torrent (index k: 1 or 2): explanatory two-level factor; three water parameters (index n: 1 to 3) measured at each site (index l: 1 to 10)[7]: water pH, water conductivity and dissolved $O_2$; fish characteristics (index o: 1 to 2), evaluated for each individual fish (index j): weight and sex; the two level interactions of fish characteristics (weight and sex) with the factor 'tailings torrent'; $(1|\text{site}_l)$: random intercept; $\varepsilon_{j|l}$: normal distributed residual, nested according to site. We adjusted separate models for each of the 12 evaluated elements. Significant effect of the tailings torrent meant that the concentration of the evaluated element in *Geophagous brasiliensis* muscle tissue differ between affected and reference sites. As far as our aim was to evaluate the effects of the tailings torrent, significant effects of water parameters or fish characteristics alone would discard our working hypothesis.

Difference in the slopes of the response variable with the explanatory variable did not test the null hypothesis that one of the slopes was equal to zero. Therefore, in order to test this null hypothesis, we resumed model simplification adjusting separate explanatory models for both 'impacted' and 'not impacted' (reference) sites. Weight was log-transformed,

7 We measured three water parameters, represented by the index n, being either 1, 2 or 3 (respectively water pH, water conductivity and dissolved O2), in each site (index k). The vertical bar (|) represents "given", meaning that water parameters are the same for all fish individuals collected in the same site.

because its effects were assumed to be multiplicative. All continuous explanatory variables were centered and scaled (using scale function in R, Becker et al. 1988), so as to avoid misestimation of their effects due to differences in their range or their values. Significance was evaluated by deletion of non-significant terms, through model comparison (using anova function in R). All models were subjected to residual analysis to evaluate adequacy. When suspected, we evaluated non-linearity by adjusting generalized additive models (GAMs). We reported exact p-values for significant terms, evaluated by deletion from the minimum model; for non-significant terms we reported the lowest p-value of them, found along model simplification. All statistical analyses were done under R (R Core Team 2018). We used the packages mgcv (Wood 2011), lme4 (Bates et al. 2015) and e1071 (Meyer et al. 2018).

### References

Bates D, Maechler M, Bolker B, and Walker S. 2015. lme4: Linear mixed-effects models using Eigen and S4. R package version 1.1–7. 2014.

Becker RA, Chambers JM. and Wilks AR. 1988. The New S Language. Wadsworth and Brooks/Cole.

Meyer, D., Dimitriadou, E., Hornik, K., Weingessel, A., & Leisch, F. (2018). e1071: Misc Functions of the Department of Statistics, Probability Theory Group (Formerly: E1071), TU Wien, 2015. R package version, 1(8).

R Core Team. 2018. R: A language and environment for statistical computing. R Foundation for Statistical Computing, Vienna, Austria. URL https://www.R-project.org/.

Wood SN. 2011. Fast stable restricted maximum likelihood and marginal likelihood estimation of semiparametric generalized linear models. *Journal of the Royal Statistical Society* (B) 73(1):3-36
