## Supplementary material for "Impacts of the Samarco tailing dam collapse on metals and arsenic concentration in freshwater fish muscle from Doce River, southeastern Brazil": SD Table S1: Fish species, number of individuals, sites and indication for which statistical analyses they were included

**Supplemental Data Table S1.** Number of fish individuals collected per species in sites that were unaffected (n=7) and affected (n=5) by the Samarco iron ore tailings dam rupture in the Doce river basin, ordered according to decreasing accumulated abundance, and number of sites where the species occurred. The last five columns point the species included in each of the statistical analyses. The numbers between parentheses refer to the equation of the statistical model mentioned in the Statistical Analyses section of the Methodology.

| sp # | Fish species | Unaffected sites | Affected sites | Total | Number of sites | Two-way MANOVA (eq. 1) | Three-way MANOVA (eq. 2) | Strict three-way MANOVA (eq. 2) | Intraspecific MANOVAs (eq. 2, 3) | GLMMs ANOVAs and ANCOVAs (eq. 4, 5) | Intraspecific GLMM ANCOVA (eq. 6) |
|---|---|---|---|---|---|---|---|---|---|---|---|
| 1 | *Geophagus brasiliensis* | 30 | 6 | 36 | 9 | x | x | x | x | x | x |
| 2 | *Oligosarcus argenteus* | 27 | 1 | 28 | 8 | x | x | | x | x | |
| 3 | *Astyanax lacustris* | 21 | 0 | 21 | 6 | x | x | | | x | |
| 4 | *Hoplias intermedius* | 10 | 7 | 17 | 8 | x | x | x | x | x | |
| 5 | *Hypostomus affinis* | 13 | 4 | 17 | 6 | x | x | x | x | x | |
| 6 | *Pachyurus adspersus* | 16 | 1 | 17 | 5 | x | x | | x | x | |
| 7 | *Loricariichthys castaneus* | 14 | 2 | 16 | 4 | x | x | | x | x | |
| 8 | *Megaleporinus conirostris* | 9 | 4 | 13 | 6 | x | x | x | | x | |
| 9 | *Astyanax* sp. 2 | 10 | 0 | 10 | 2 | x | x | | | x | |
| 10 | *Hypomasticus mormyrops* | 10 | 0 | 10 | 2 | x | x | | | x | |
| 11 | *Pimelodus maculatus* | 3 | 4 | 7 | 3 | x | x | x | | x | |
| 12 | *Oreochromis niloticus* | 4 | 3 | 7 | 4 | x | x | x | | x | |
| 13 | *Henochilus wheatlandii* | 7 | 0 | 7 | 2 | x | x | | | x | |
| 14 | *Hoplias malabaricus* | 1 | 5 | 6 | 2 | x | x | | | x | |
| 15 | *Rhamdia quelen* | 2 | 3 | 5 | 3 | x | x | | | x | |
| 16 | *Hypostomus auroguttatus* | 5 | 0 | 5 | 4 | x | x | | | x | |
| 17 | *Cichla piquiti* | 4 | 0 | 4 | 1 | x | x | | | x | |
| 18 | *Leporinus copelandii* | 4 | 0 | 4 | 3 | x | x | | | x | |
| 19 | *Salminus brasiliensis* | 0 | 3 | 3 | 1 | x | x | | | x | |
| 20 | *Prochilodus vimboides* | 0 | 2 | 2 | 1 | x | x | | | x | |
| 21 | *Lophiosilurus alexandri* | 1 | 1 | 2 | 2 | x | x | | | x | |
| 22 | *Trachelyopterus striatulus* | 1 | 1 | 2 | 1 | x | x | | | x | |
| 23 | *Astyanax fasciatus* | 2 | 0 | 2 | 2 | x | x | | | x | |
| 24 | *Crenicichla lacustris* | 2 | 0 | 2 | 2 | x | x | | | x | |
| 25 | *Cyphocharax gilbert* | 2 | 0 | 2 | 1 | x | x | | | x | |
| 26 | *Delturus carinotus* | 2 | 0 | 2 | 2 | x | x | | | x | |
| 27 | *Astyanax* sp. 1 | 0 | 1 | 1 | 1 | x | x | | | x | |
| 28 | *Hoplosternum littorale* | 0 | 1 | 1 | 1 | x | x | | | x | |
| 29 | *Leporinus macrocephalus* | 0 | 1 | 1 | 1 | x | x | | | x | |
| 30 | *Myloplus asterias* | 0 | 1 | 1 | 1 | x | x | | | x | |
| 31 | *Prochilodus costatus* | 0 | 1 | 1 | 1 | x | x | | | x | |
| 32 | *Prochilodus lineatus* | 0 | 1 | 1 | 1 | x | x | | | x | |
| 33 | *Gymnotus carapo* | 1 | 0 | 1 | 1 | x | x | | | x | |
| 34 | *Pseudauchenipterus affinis* | 1 | 0 | 1 | 1 | x | x | | | x | |
| | **Total** | **202** | **53** | **255** | **12** | **34** | **34** | **6** | **6** | **34** | **1** |
