## Supplementary material for "Impacts of the Samarco tailing dam collapse on metals and arsenic concentration in freshwater fish muscle from Doce River, southeastern Brazil": SD Table S2: Results of the two-way MANOVA for all fish species

**Supplemental Data Table S2.** Effects of the tailings mudflow that resulted from the Samarco iron ore dam break on the profile of elements' concentration in fish muscle tissue. Results of the two-way MANOVA for all fish species (eq. 1 in the Supplemental Data Methodology Details).

| Term | d.f. | Pillai | aprox. F | num. d.f. | den. d.f. | P (>F) | Significance |
|---|---|---|---|---|---|---|---|
| Tailing mudflow | 1 | 0.40698 | 13.2682 | 12 | 232 | < 2.2e-16 | *** |
| Site (block) | 10 | 1.60912 | 3.8514 | 120 | 2410 | < 2.2e-16 | *** |
| Residuals | 243 | | | | | | |
