## Supplementary material for "Impacts of the Samarco tailing dam collapse on metals and arsenic concentration in freshwater fish muscle from Doce River, southeastern Brazil": SD Table S3: Results of the three-way MANOVA for all fish species

**Supplementary Data Table S3.** Effects of the tailings mudflow that resulted from the Samarco iron ore dam break, of the fish species and its interaction, on the profile of elements' concentration in fish muscle tissue. Results of the three-way MANOVA for all fish species (eq. 2 in the Supplemental Data Methodology Details).

| Term | d.f. | Pillai | aprox. F | num. d.f. | den. d.f. | P (>F) | Significance |
|---|---|---|---|---|---|---|---|
| Tailing mudflow | 1 | 0.6499 | 29.085 | 12 | 188 | < 2e-16 | *** |
| Fish species | 33 | 4.4018 | 3.4935 | 396 | 2388 | < 2e-16 | *** |
| Tailing:species | 12 | 0.8639 | 1.2865 | 108 | 2388 | 0.01426 | * |
| Site (block) | 9 | 1.6109 | 3.5608 | 144 | 1764 | < 2e-16 | *** |
| Residuals | 199 | | | | | | |
