## Supplementary material for "Impacts of the Samarco tailing dam collapse on metals and arsenic concentration in freshwater fish muscle from Doce River, southeastern Brazil": SD Table S4: Results of the two-way MANOVA for the six fish species that presented at least three individuals in each affected and reference sites

**Supplemental Data Table S4.** Effects of the tailings mudflow that resulted from the Samarco iron ore dam break on the profile of elements' concentration in fish muscle tissue. Results of the two-way MANOVA for the six fish species that presented at least three individuals in each affected and reference sites (n=97 individuals; eq. 2 in the Supplemental Data Methodology Details).

| Term | d.f. | Pillai | aprox. F | num. d.f. | den. d.f. | P (>F) | Significance |
|---|---|---|---|---|---|---|---|
| Tailing mudflow | 1 | 0.61437 | 8.7623 | 12 | 66 | 1.07E-09 | *** |
| Fish species | 5 | 2.18516 | 4.5284 | 60 | 350 | < 2e-16 | *** |
| Tailing:species | 8 | 1.02949 | 1.5125 | 96 | 584 | 0.01256 | * |
| Site (block) | 5 | 2.10091 | 2.1665 | 60 | 350 | 2.41E-08 | *** |
| Residuals | 77 | | | | | | |
