## Supplementary material for "Impacts of the Samarco tailing dam collapse on metals and arsenic concentration in freshwater fish muscle from Doce River, southeastern Brazil": SD Table S5: Results of the two or three-way MANOVAs

**Supplemental Data Table S5.** Effects of the tailings torrent that resulted from the Samarco iron ore dam break on the profile of elements' concentration in fish muscle tissue, within the six more abundant species. Results of the two or three-way MANOVAs (eqs. 1 or 3 in the Supplemental Data Methodology Details).

| Species | Term | d.f. | Pillai | aprox. F | num. d.f. | den. d.f. | P (>F) | Significance |
|---|---|---|---|---|---|---|---|---|
| *Geophagus brasiliensis* | | | | | | | | |
| | Tailings mudflow | 1 | 0.842 | 7.1275 | 12 | 16 | 2.31E-04 | *** |
| | Site (block) | 7 | 3.64 | 1.9859 | 84 | 154 | 1.18E-04 | *** |
| | Residuals | 27 | | | | | | |
| *Oligosarcus argenteus* | | | | | | | | |
| | Tailings mudflow | 1 | 0.659 | 1.4501 | 12 | 9 | 0.29284 | |
| | Site (block) | 6 | 1.607 | 1.607 | 72 | 84 | 0.01828 | * |
| | Residuals | 20 | | | | | | |
| *Hoplias intermedius* | | | | | | | | |
| | Tailings mudflow | 1 | 0.788 | 1.2358 | 12 | 4 | 0.4572 | |
| | Residuals | 15 | | | | | | |
| *Hypostomus affinis* | | | | | | | | |
| | Tailings mudflow | 1 | 0.864 | 2.1232 | 12 | 4 | 0.2437 | |
| | Residuals | 15 | | | | | | |
| *Pachyurus adspersus* | | | | | | | | |
| | Tailings mudflow | 1 | 0.994 | 57.55 | 12 | 4 | 0.000683 | *** |
| | Residuals | 15 | | | | | | |
| *Loricariichthys castaneus* | | | | | | | | |
| | Tailings mudflow | 1 | 0.978 | 3.7535 | 12 | 1 | 0.3849 | |
| | Site (block) | 2 | 1.913 | 3.6483 | 24 | 4 | 0.1082 | |
| | Residuals | 12 | | | | | | |
