## Supplementary material for "Impacts of the Samarco tailing dam collapse on metals and arsenic concentration in freshwater fish muscle from Doce River, southeastern Brazil": SD Table S6: Mean concentration of metals and arsenic in fish from sites unaffected and affected and results of the mixed effects one-way ANOVAs

**Supplementary Data Table S6.** Mean concentration in fish muscle tissue from sites unaffected and affected by the Samarco iron ore tailings dam rupture, and results of the mixed effects one-way ANOVAs (GLMMs, eq. 4 in the Supplemental Data Methodology Details) to evaluate the effect of the tailings mudflow that resulted from the iron ore dam brake on the concentration of elements in fish muscle tissue. Site was adjusted as random intercept.

| Element | Unaffected | Affected | LRT | P value | Significance |
|---|---|---|---|---|---|
| Ag | 0.00066 | 0.00037 | 4.4016 | 0.03591 | * |
| Al | 14.25 | 11.13 | 0.0269 | 0.8698 | n.s. |
| As | 0.031 | 0.11 | 14.0190 | 0.000181 | *** |
| Cd | 0.00068 | 0.00060 | 0.5597 | 0.4544 | n.s. |
| Cr | 0.086 | 0.021 | 0.2968 | 0.5859 | n.s. |
| Cu | 0.28 | 0.25 | 0.0673 | 0.7138 | n.s. |
| Fe | 12.93 | 15.14 | 2.1205 | 0.1453 | n.s. |
| Hg | 0.103 | 0.20 | 5.7268 | 0.01671 | * |
| Mn | 1.405 | 1.23 | 0.0217 | 0.8828 | n.s. |
| Ni | 0.025 | 0.027 | 0.5102 | 0.4751 | n.s. |
| Pb | 0.013 | 0.0081 | 0.0761 | 0.7827 | n.s. |
| Zn | 11.92 | 9.68 | 4.1503 | 0.04163 | * |
