## Supplementary material for "Impacts of the Samarco tailing dam collapse on metals and arsenic concentration in freshwater fish muscle from Doce River, southeastern Brazil": SD Table S7: Results of the mixed effects ANCOVAs

**Supplemental Data Table S7.** Effects of the tailings mudflow that resulted from the Samarco iron ore dam break on the concentration of elements in fish muscle tissue, with water parameters and fish characteristics as co-variables. Results of the mixed effects ANCOVAs (GLMMs, eq. 6 in the Supplemental Data Methodology Details). Site was adjusted as random intercept. Significance was evaluated by deletion of non-significant terms. LRT: Likelyhood ration test; P(>Chi): p-value; n.s.: P > 0.05; *: P<0.05; **: P < 0.01; ***: P < 0.001. (1) When a term was included in a significant interaction, it could not be tested separately (Crawley, 2013).

| Element | Terms | d.f. | LRT | P(>Chi) | Significance |
|---|---|---|---|---|---|
| **Ag** | Tailing mudflow | 1 | 0.053 | 0.8179 | n.s. |
| | Fish sex | 1 | 0.651 | 0.4198 | n.s. |
| | Fish weight | 1 | 5.759 | 0.0164 | * |
| | Water pH | 1 | 7.429 | 0.0064 | ** |
| | Water $O_2$ | 1 | 7.687 | 0.0056 | *** |
| | Water conductivity | 1 | 1.168 | 0.2798 | n.s. |
| | mudflow*Sex | 1 | 2.431 | 0.1189 | n.s. |
| | mudflow*Weight | 1 | 0.824 | 0.3640 | n.s. |
| **Al** | Tailing mudflow | 1 | 5.639 | 0.0176 | * |
| | Fish sex | 1 | 0.429 | 0.5123 | n.s. |
| | Fish weight | 1 | 3.453 | 0.0631 | n.s. |
| | Water pH | 1 | 0.002 | 0.9650 | n.s. |
| | Water $O_2$ | 1 | 0.219 | 0.6396 | n.s. |
| | Water conductivity | 1 | 8.633 | 0.0033 | ** |
| | mudflow*Sex | 1 | 1.869 | 0.1717 | n.s. |
| | mudflow*Weight | 1 | 1.420 | 0.2335 | n.s. |
| **As** | Tailing mudflow | 1 | - | - | (1) |
| | Fish sex | 1 | - | - | (1) |
| | Fish weight | 1 | - | - | (1) |
| | Water pH | 1 | 1.703 | 0.1919 | n.s. |
| | Water $O_2$ | 1 | 0.046 | 0.8302 | n.s. |
| | Water conductivity | 1 | 3.380 | 0.0660 | n.s. |
| | mudflow*Sex | 1 | 5.590 | 0.0181 | * |
| | mudflow*Weight | 1 | 21.418 | 3.69E-06 | *** |
| **Cd** | Tailing mudflow | 1 | 0.321 | 0.5712 | n.s. |
| | Fish sex | 1 | 0.021 | 0.8847 | n.s. |
| | Fish weight | 1 | 2.743 | 0.0977 | n.s. |
| | Water pH | 1 | 2.119 | 0.1455 | n.s. |
| | Water $O_2$ | 1 | 3.021 | 0.0822 | n.s. |
| | Water conductivity | 1 | 1.225 | 0.2685 | n.s. |
| | mudflow*Sex | 1 | 0.035 | 0.8513 | n.s. |
| | mudflow*Weight | 1 | 0.776 | 0.3785 | n.s. |
| **Cr** | Tailing mudflow | 1 | 0.044 | 0.8341 | n.s. |
| | Fish sex | 1 | 0.220 | 0.6391 | n.s. |
| | Fish weight | 1 | 5.220 | 0.0223 | * |
| | Water pH | 1 | 0.037 | 0.8485 | n.s. |
| | Water $O_2$ | 1 | 0.749 | 0.3869 | n.s. |
| | Water conductivity | 1 | 2.478 | 0.1155 | n.s. |
| | mudflow*Sex | 1 | 2.734 | 0.0983 | n.s. |
| | mudflow*Weight | 1 | 0.575 | 0.4482 | n.s. |

**Supplemental Data Table S7.** *Continuation*

| Element | Terms | d.f. | LRT | P(>Chi) | Significance |
|---|---|---|---|---|---|
| **Cu** | Tailing mudflow | 1 | - | - | (1) |
| | Fish sex | 1 | 8.717 | 0.0032 | ** |
| | Fish weight | 1 | - | - | (1) |
| | Water pH | 1 | 0.812 | 0.3676 | n.s. |
| | Water $O_2$ | 1 | 1.261 | 0.2615 | n.s. |
| | Water conductivity | 1 | 1.725 | 0.1890 | n.s. |
| | mudflow*Sex | 1 | 0.694 | 0.4049 | n.s. |
| | mudflow*Weight | 1 | 4.035 | 0.0446 | * |
| **Fe** | Tailing mudflow | 1 | 0.401 | 0.5268 | n.s. |
| | Fish sex | 1 | 2.093 | 0.1480 | n.s. |
| | Fish weight | 1 | 0.013 | 0.9103 | n.s. |
| | Water pH | 1 | 0.149 | 0.6991 | n.s. |
| | Water $O_2$ | 1 | 0.500 | 0.4793 | n.s. |
| | Water conductivity | 1 | 3.439 | 0.0637 | n.s. |
| | mudflow*Sex | 1 | 2.557 | 0.1098 | n.s. |
| | mudflow*Weight | 1 | 1.788 | 0.1812 | n.s. |
| **Hg** | Tailing mudflow | 1 | 10.462 | 0.0012 | ** |
| | Fish sex | 1 | 5.763 | 0.0164 | * |
| | Fish weight | 1 | 0.357 | 0.5502 | n.s. |
| | Water pH | 1 | 0.897 | 0.3437 | n.s. |
| | Water $O_2$ | 1 | 1.722 | 0.1894 | n.s. |
| | Water conductivity | 1 | 7.784 | 0.0053 | ** |
| | mudflow*Sex | 1 | 1.242 | 0.2651 | n.s. |
| | mudflow*Weight | 1 | 0.293 | 0.5884 | n.s. |
| **Mn** | Tailing mudflow | 1 | 0.948 | 0.3303 | n.s. |
| | Fish sex | 1 | 0.311 | 0.5769 | n.s. |
| | Fish weight | 1 | 11.532 | 0.0007 | *** |
| | Water pH | 1 | 0.670 | 0.4129 | n.s. |
| | Water $O_2$ | 1 | 3.345 | 0.0674 | n.s. |
| | Water conductivity | 1 | 0.376 | 0.5398 | n.s. |
| | mudflow*Sex | 1 | 1.555 | 0.2124 | n.s. |
| | mudflow*Weight | 1 | 2.670 | 0.1023 | n.s. |
| **Ni** | Tailing mudflow | 1 | 0.016 | 0.9002 | n.s. |
| | Fish sex | 1 | 0.964 | 0.3261 | n.s. |
| | Fish weight | 1 | 0.397 | 0.5289 | n.s. |
| | Water pH | 1 | 1.859 | 0.1727 | n.s. |
| | Water $O_2$ | 1 | 3.012 | 0.0826 | n.s. |
| | Water conductivity | 1 | 4.064 | 0.0438 | * |
| | mudflow*Sex | 1 | 0.147 | 0.7011 | n.s. |
| | mudflow*Weight | 1 | 0.474 | 0.4914 | n.s. |

*continues*

**Supplemental Data Table S7.** *Continuation*

| Element | Terms | d.f. | LRT | P(>Chi) | Significance |
|---|---|---|---|---|---|
| **Pb** | Tailing mudflow | 1 | 2.173 | 0.1405 | n.s. |
| | Fish sex | 1 | 1.802 | 0.1795 | n.s. |
| | Fish weight | 1 | 17.016 | 3.71E-05 | *** |
| | Water pH | 1 | 1.252 | 0.2633 | n.s. |
| | Water $O_2$ | 1 | 1.370 | 0.2418 | n.s. |
| | Water conductivity | 1 | 6.206 | 0.0127 | * |
| | mudflow*Sex | 1 | 1.276 | 0.2587 | n.s. |
| | mudflow*Weight | 1 | 2.696 | 0.1006 | n.s. |
| **Zn** | Tailing mudflow | 1 | - | - | (1) |
| | Fish sex | 1 | 0.035 | 0.8525 | n.s. |
| | Fish weight | 1 | - | - | (1) |
| | Water pH | 1 | 0.001 | 0.9775 | n.s. |
| | Water $O_2$ | 1 | 0.002 | 0.9672 | n.s. |
| | Water conductivity | 1 | 0.586 | 0.4441 | n.s. |
| | mudflow*Sex | 1 | 0.910 | 0.3399 | n.s. |
| | mudflow*Weight | 1 | 4.629 | 0.0314 | * |
