## Supplementary material for "Impacts of the Samarco tailing dam collapse on metals and arsenic concentration in freshwater fish muscle from Doce River, southeastern Brazil": SD Table S8: Results of the mixed effects ANCOVAs for Geophagous brasiliensis

**Supplementary Data Table S8.** Intraspecific effects of the tailings mudflow that resulted from the Samarco iron ore dam break on the concentration of elements in with water parameters and fish characteristics as co-variables on elements' concentration in *Geophagous brasiliensis* muscle tissue. Results of the mixed effects ANCOVAs (GLMMs, eq. 7 in the Supplemental Data Methodology Details). Site was adjusted as random intercept. Significance was evaluated by deletion of non-significant terms. LRT: Likelyhood ration test; P(>Chi): p-value; n.s.: P > 0.05; *: P<0.05; **: P < 0.01; ***: P < 0.001. (1) When a term was included in a significant interaction, it could not be tested separately (Crawley, 2013).

| **Element** | **Term** | **d.f.** | **LRT** | **P(>Chi)** | **Significance** |
|---|---|---|---|---|---|
| **Ag** | Tailing mudflow | 1 | 1.803 | 0.1794 | n.s. |
| | Fish sex | 1 | 0.071 | 0.7906 | n.s. |
| | Fish weight | 1 | 2.340 | 0.1261 | n.s. |
| | Water pH | 1 | 12.012 | 0.00053 | *** |
| | Water $O_2$ | 1 | 3.127 | 0.0770 | n.s. |
| | Water conductivity | 1 | <0.001 | 0.9947 | n.s. |
| **Al** | Tailing mudflow | 1 | 1.020 | 0.3124 | n.s. |
| | Fish sex | 1 | 0.224 | 0.6363 | n.s. |
| | Fish weight | 1 | 0.319 | 0.5720 | n.s. |
| | Water pH | 1 | 0.226 | 0.6344 | n.s. |
| | Water $O_2$ | 1 | 0.038 | 0.8461 | n.s. |
| | Water conductivity | 1 | 2.030 | 0.1542 | n.s. |
| **As** | Tailing mudflow | 1 | 11.187 | 8.24E-04 | *** |
| | Fish sex | 1 | 0.219 | 0.6401 | n.s. |
| | Fish weight | 1 | 7.090 | 0.0078 | ** |
| | Water pH | 1 | 0.006 | 0.9383 | n.s. |
| | Water $O_2$ | 1 | 0.167 | 0.6829 | n.s. |
| | Water conductivity | 1 | 0.872 | 0.3505 | n.s. |
| **Cd** | Tailing mudflow | 1 | 2.548 | 0.1104 | n.s. |
| | Fish sex | 1 | 1.890 | 0.1692 | n.s. |
| | Fish weight | 1 | 3.155 | 0.0757 | n.s. |
| | Water pH | 1 | 3.290 | 0.0697 | n.s. |
| | Water $O_2$ | 1 | 4.020 | 0.0450 | * |
| | Water conductivity | 1 | 0.459 | 0.4979 | n.s. |
| **Cr** | Tailing mudflow | 1 | 1.368 | 0.2422 | n.s. |
| | Fish sex | 1 | 0.036 | 0.8486 | n.s. |
| | Fish weight | 1 | 0.048 | 0.8274 | n.s. |
| | Water pH | 1 | 0.171 | 0.6789 | n.s. |
| | Water $O_2$ | 1 | 0.025 | 0.8752 | n.s. |
| | Water conductivity | 1 | 0.551 | 0.4581 | n.s. |
| **Cu** | Tailing mudflow | 1 | 2.249 | 0.1337 | n.s. |
| | Fish sex | 1 | 0.118 | 0.7312 | n.s. |
| | Fish weight | 1 | 6.211 | 0.0127 | * |
| | Water pH | 1 | 2.036 | 0.1537 | n.s. |
| | Water $O_2$ | 1 | 4.489 | 0.0341 | * |
| | Water conductivity | 1 | 0.524 | 0.4693 | n.s. |

*continues*

**Supplementary Data Table S8.** *Continuation*

| Element | Term | d.f. | LRT | P(>Chi) | Significance |
|---|---|---|---|---|---|
| **Fe** | Tailing mudflow | 1 | 1.893 | 0.1689 | n.s. |
| | Fish sex | 1 | 0.033 | 0.8556 | n.s. |
| | Fish weight | 1 | 0.064 | 0.8005 | n.s. |
| | Water pH | 1 | 0.047 | 0.8291 | n.s. |
| | Water $O_2$ | 1 | 0.677 | 0.4107 | n.s. |
| | Water conductivity | 1 | 2.879 | 0.0897 | n.s. |
| **Hg** | Tailing mudflow | 1 | 15.482 | 8.33E-05 | *** |
| | Fish sex | 1 | 0.090 | 0.1134 | n.s. |
| | Fish weight | 1 | 0.987 | 0.3205 | n.s. |
| | Water pH | 1 | 0.013 | 0.9084 | n.s. |
| | Water $O_2$ | 1 | 0.150 | 0.6987 | n.s. |
| | Water conductivity | 1 | 10.021 | 0.0015 | ** |
| **Mn** | Tailing mudflow | 1 | 0.017 | 0.8957 | n.s. |
| | Fish sex | 1 | 0.749 | 0.3867 | n.s. |
| | Fish weight | 1 | 7.650 | 0.0057 | *** |
| | Water pH | 1 | 2.860 | 0.0908 | n.s. |
| | Water $O_2$ | 1 | 0.013 | 0.9107 | n.s. |
| | Water conductivity | 1 | 0.200 | 0.6547 | n.s. |
| **Ni** | Tailing mudflow | 1 | 0.034 | 0.8529 | n.s. |
| | Fish sex | 1 | 2.981 | 0.0842 | n.s. |
| | Fish weight | 1 | 7.786 | 0.0053 | ** |
| | Water pH | 1 | 0.282 | 0.5955 | n.s. |
| | Water $O_2$ | 1 | 0.495 | 0.4817 | n.s. |
| | Water conductivity | 1 | 2.584 | 0.1080 | n.s. |
| **Pb** | Tailing mudflow | 1 | 0.117 | 0.7324 | n.s. |
| | Fish sex | 1 | 0.355 | 0.5514 | n.s. |
| | Fish weight | 1 | 1.643 | 0.1999 | n.s. |
| | Water pH | 1 | 2.243 | 0.1342 | n.s. |
| | Water $O_2$ | 1 | 1.409 | 0.2352 | n.s. |
| | Water conductivity | 1 | 6.756 | 0.0093 | *** |
| **Zn** | Tailing mudflow | 1 | 1.882 | 0.1701 | n.s. |
| | Fish sex | 1 | 1.222 | 0.2690 | n.s. |
| | Fish weight | 1 | 4.526 | 0.0334 | * |
| | Water pH | 1 | 0.799 | 0.3715 | n.s. |
| | Water $O_2$ | 1 | 0.570 | 0.4501 | n.s. |
| | Water conductivity | 1 | 0.170 | 0.6801 | n.s. |
