## Supplementary material for "Impacts of the Samarco tailing dam collapse on metals and arsenic concentration in freshwater fish muscle from Doce River, southeastern Brazil": SD Table S9: Mean + standard deviation (SD) of water parameters

**Supplementary Data Table S9.** Mean ± standard deviation (SD) of water parameters in sites unaffected (n=7) and affected (n=5) by the Samarco iron ore tailings dam rupture.

| Water parameter | Unaffected | | Affected | |
|---|---|---|---|---|
| | Mean | SD | Mean | SD |
| pH | 7.89 | 0.27 | 8.05 | 0.11 |
| Dissolved $O_2$ ($mg.L^{-1}$) | 99.85 | 11.28 | 94.53 | 6.86 |
| Conductivity ($\mu S.cm^{-1}$) | 45.93 | 18.27 | 74.50 | 13.67 |
